# Tracking the circulating SARS-CoV-2 variants in Turkey: Complete genome sequencing and molecular characterization of 1000 SARS-CoV-2 samples

**DOI:** 10.1101/2022.04.19.488722

**Authors:** Faruk Berat Akçeşme, Tuğba Kul Köprülü, Burçin Erkal, Şeyma İş, Birsen Cevher Keskin, Betül Akçeşme, Kürşad Nuri Baydili, Bahar Gezer, Jülide Balkan, Bihter Uçar, Osman Gürsoy, Mehmet Taha Yıldız, Halil Kurt, Nevzat Ünal, Mustafa Altındiş, Celalettin Korkmaz, Hasan Türkez, Özlem Bayraktar, Barış Demirkol, Yasemin Çağ, Melih Akay Arslan, Hilal Abakay, Şükran Köse, Abdülkadir Özel, Neslihan Mutluay, Şaban Tekin

## Abstract

Severe acute respiratory syndrome coronavirus 2 (SARS-CoV-2) is a highly transmissible coronavirus and has caused a pandemic of acute respiratory disease, named ‘coronavirus disease 2019’ (COVID-19). COVID-19 has a deep impact on public health as one of the most serious pandemics in the last century. Tracking SARS-CoV-2 is important for monitoring and assessing its evolution. This is only possible by detecting all mutations in the viral genome through genomic sequencing. Moreover, accurate detection of SARS-CoV-2 and tracking its mutations is also required for its correct diagnosis. Potential effects of mutations on the prognosis of the disease can be observed. Assignment of epidemiological lineages in an emerging pandemic requires efforts. To address this, we collected 1000 SARS-CoV-2 samples from different geographical regions in Turkey and analyze their genome comprehensively. To track the virus across Turkey we focus on 10 distinct cities in different geographic regions. Each SARS-CoV-2 genome was analyzed and named according to the nomenclature system of Nextclade and Pangolin Lineage. Furthermore, the frequency of the variations observed in 10 months was also determined by region. In this way, we have observed how the virus mutations and what kind of transmission mechanism it has. The effects of age and disease severity on lineage distribution were other considered parameters. The temporal rates of SARS-CoV-2 variants by time in Turkey were close to the global trend. This study is one of the most comprehensive whole genome analyses of SARS-CoV-2 that represents a general picture of the distribution of SARS-CoV-2 variations in Turkey in 2021.

**Author Summary:** Since the outbreak of the COVID-19 pandemic in 2019, the viral genome of SARS-CoV-2 was analysed intensively all over the world both to detect its zoonotic origin and the emerging variants worldwide together with the variants’ effect on the prognosis and treatment, respectively, of the infection. Remarkable COVID-19 studies were also made in Turkey as it was in the rest of the world. To date, indeed, almost all studies on COVID-19 in Turkey either sequenced only a small number of the viral genome or analysed the viral genome which was obtained from online databases. In respect thereof, our study constitutes a milestone regarding both the huge sample size consisting of 1000 viral genomes and the widespread geographic origin of the viral genome samples. Our study provides new insights both into the SARS-CoV-2 landscape of Turkey and the transmission of the emerging viral pathogen and its interaction with its vertebrate host.

## Introduction

COVID-19, the worst known pandemic of the last century, remains a serious threat to public health (1). Numerous scientific studies have been carried out regarding the virus, which has been on the world agenda since the day it emerged in Wuhan, China, on 31 December 2019 (2). The coronavirus family, well known by the scientific world, has a new virus member that has the characteristic of rapid human to-human transmission. The first analyzed genome of SARS-CoV-2, which is generally accepted to be transmitted from animal to human, has undergone many mutations so far.

Observing these mutations, investigating the effects of these mutations on the prognosis of the disease and the rate of transmission became the primary targets of scientific research (3). Despite this, it was not straightforward to detect all mutations of the viral genome during the fight against the pandemic and to determine how these mutations affect the prognosis of the disease since diagnostic tools do not provide this information (4).

While this struggle continues in scientific circles, decision-makers and international authorities have tried to take many measures, from isolation to restrictions, to reduce the speed of the pandemic. The mutations in the virus began to affect people’s working, social and economic life directly and very quickly.

Despite all kinds of prevention, precautions and struggle, SARS-CoV-2 became effective all over the world and infected more than 400 million people while taking more than 6 million lives (5).

The first COVID-19 case in Turkey was announced by the Ministry of Health Turkey on 11 March 2020. To date in Turkey, there have been around 14,2 Million confirmed cases of COVID-19 with 95 thousand deaths (data from (6) (accessed: 20 March 2022).

SARS-CoV-2 is a single-stranded, enveloped RNA virus. Its genome, comprised of 29.9 kilobase pairs, has 11 coding regions, namely: ORF1ab –the longest coding region–, ORF3a, ORF6, ORF7a, ORF7b, ORF8, ORF10, spike (S), envelope (E), membrane (M) and nucleocapsid (N) (7–9).

The specificity, transmissibility, pathogenicity, and response of SARS-CoV-2 to treatments have changed day by day with variations in the virus genome. Among the mutations, the D614G mutation in the spike protein is the first to come to the fore, and studies have shown that this mutation is more infectious than the first isolate and has higher viral loads (10). In the Alpha variant (B.1.1.7 lineage), which was first detected in the United Kingdom in 2020, amino acid 69 and 70 deletion (Δ69/70) and amino acid 144 deletion (Δ144) in the N-terminal domain and N501Y mutations found in the receptor-binding domain. Although Δ69/70 is not resistant to monoclonal antibodies, it has been found to confer a competence that results in more enhanced cell-cell fusion than the D614G variant and increases the infectivity of the virus. Towards the end of 2020, 11 mutations in spike protein were identified in the Delta variant (B.1.617.2 lineage), which was first detected in India; T19R, T95I, G142D, Δ156–157 (amino acid 156 and 157 deletion), R158G, L452R, T478K, D614G, P681R, and D950N. Of these mutations, it was observed that the P681R mutation further facilitates furin cleavage and is highly conserved in this lineage, resulting in increased fusogenicity and pathogenicity, thus rapidly affecting the whole world. Finally, it was determined that the Omicron variant (B.1.1.529 lineage), which swept the world and was first seen in South Africa in 2021, had about 34 mutations only in the spike protein. Some of these mutations have been found to be more infectious than the Delta variant. For example, N501Y increases binding to the ACE2 receptor, which can increase transmission, and the combination of N501Y and Q498R can further increase binding affinity (11). N439K increases the affinity for ACE2, too (12). ORF1ab P4715L and S protein D614G variants were found to be linked to higher fatality rates by affecting the presentation to MHC-I and MHC-II which in turn influences the extent of the immune response. Furthermore, D614G was found to be associated with higher viral loads and interacts with ACE2 (13,14). Another mutation, namely E484K of spike RBD, allows the virus to escape from antibodies (12). In addition, Gunadi et al. showed that patients with multiple S protein mutations are about five times more likely to be hospitalized than patients with no or a single S protein mutation (13). As can be seen from the examples above, different virus variants can result in different disease severity and hospitalization risk.

To develop drugs and vaccines, genetic differences and functions of these coding regions of SARS-CoV-2 must be thoroughly researched (15). So far, many genome analyzes have been conducted to study to evolution of SARS-CoV-2 (16–20). However, very extensive studies concerning global genetic diversity and conserved regions of the viral genome must be carried out, not only to develop epitope-based vaccines, but also to develop kits for RT-PCR and thus to be able to identify the virus in a much more specific manner (21).

Different classification systems have been proposed by several research groups such as GISAID (22),_World Health Organization (23), Nextclade (24), and PANGO lineages (25), to track the evolution of SARS-CoV-2 over time, since it has natural expanding genetic diversity.

Two classification systems used in this research are PANGO lineages and Nextclade. Phylogenetic Assignment of Named Global Outbreak Lineages (Pangolin) is a dynamic virus nomenclature system, based on phylogenetic information analysis that helps to understand and to track the patterns and determinants of the spread of SARS-CoV-2 globally (25). Nextclade is another popular nomenclature system, which aims to label genetically well-defined clades when mutations reach significant frequency (20%) and global geographic spread. Names of the clades are designed as memorable and informative enough, consisting of the year it arose and a letter (24).

SARS-CoV-2 genome analyses with patient information (demographic) in Turkey has done with small number of viral genomes (26–30). Our main objective was to perform whole-genome sequencing of 1000 samples collected from different regions of Turkey in order to characterize SARS-CoV-2 genomes. Here, we identified the lineages and clades of complete genome sequences of SARS-CoV-2. With this information, we tracked the spread of the virus and assisted the genomic epidemiology of SARS-CoV-2 in Turkey. Our study is the first and the most comprehensive SARS-CoV-2 genome analysis in which collected more than 1000 viral samples traced in single research in Turkey during the COVID-19 pandemics.

## Results and Discussion

The virus that causes COVID-19, SARS-CoV-2, was changed over time like all viruses. Most changes in the viral genome have little to no impact on the virus’ properties. However, viral genomes changes may affect how easily it spreads and the associated disease severity. Furthermore, the diagnostic tools and performance of vaccines may differ because of the changes in the virus.

SARS-CoV-2 genome studies are conducted in many countries to give an effective direction to the detection, prognosis, and treatment of the disease, especially by revealing the genome of the virus and finding possible variations. In this study, collected vRNA samples taken from patients in different geographical regions of Turkey were sequenced and subjected to bioinformatics analysis. The spread of the virus pattern was identified. Each of the variants of concerns (VOC) and variant of interest (VOI) distributions was detected. We first determined each of the viral genome’s clades in all samples. As seen in the graph, the most seen clade in Turkey is 20A between March 2021 and December 2021.

We also classified the genomes according to the Pango Lineage nomenclature system. As it is seen in the table one of the most seen lineages is B.1.617.2 which is circulating mostly in India and later in Europe and USA (19). It comprised around 20% of sequences from some regions and contains L452R and E484Q mutations.

B.1.1.7. UK lineage of concern, associated with the N501Y mutation is one of the other lineages that is seen with high frequency. As it is seen in the table below, in the time interval that we collected samples there are many variants of concerns and variant of the interests.

We tried to monitor the circulation of variations in a Turkey as a country with high genetic heterogeneity and high population mobility. The table below shows the variation distributions of SARS-COV-2 according to the Nextclade nomenclature system of the samples taken from various cities. Sample collection time periods were considered, and samples are taken from the cities in parts, not in one go in different time interval in a 9-month period. As seen in the table below, there are some variations in distribution of the clades among the cities.

In Figure 4, we showed the variation distributions of SARS-CoV-2 variations according to the Pango Lineage nomenclature system of the samples taken from various cities.

**Fig 1.**
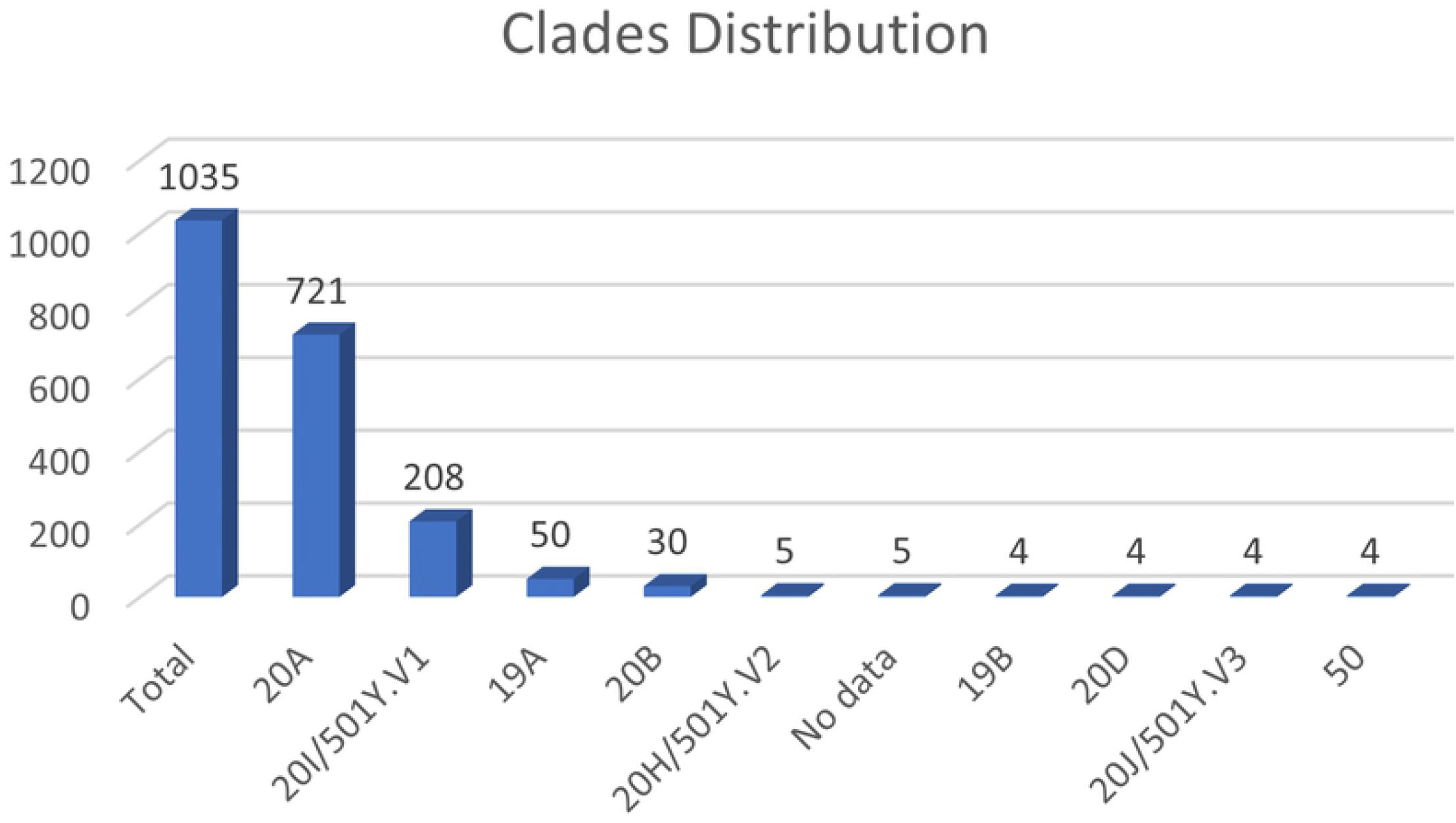
Nextclade Clades distributions of the SARS-CoV-2 distributions in Turkey.

**Fig 2.**
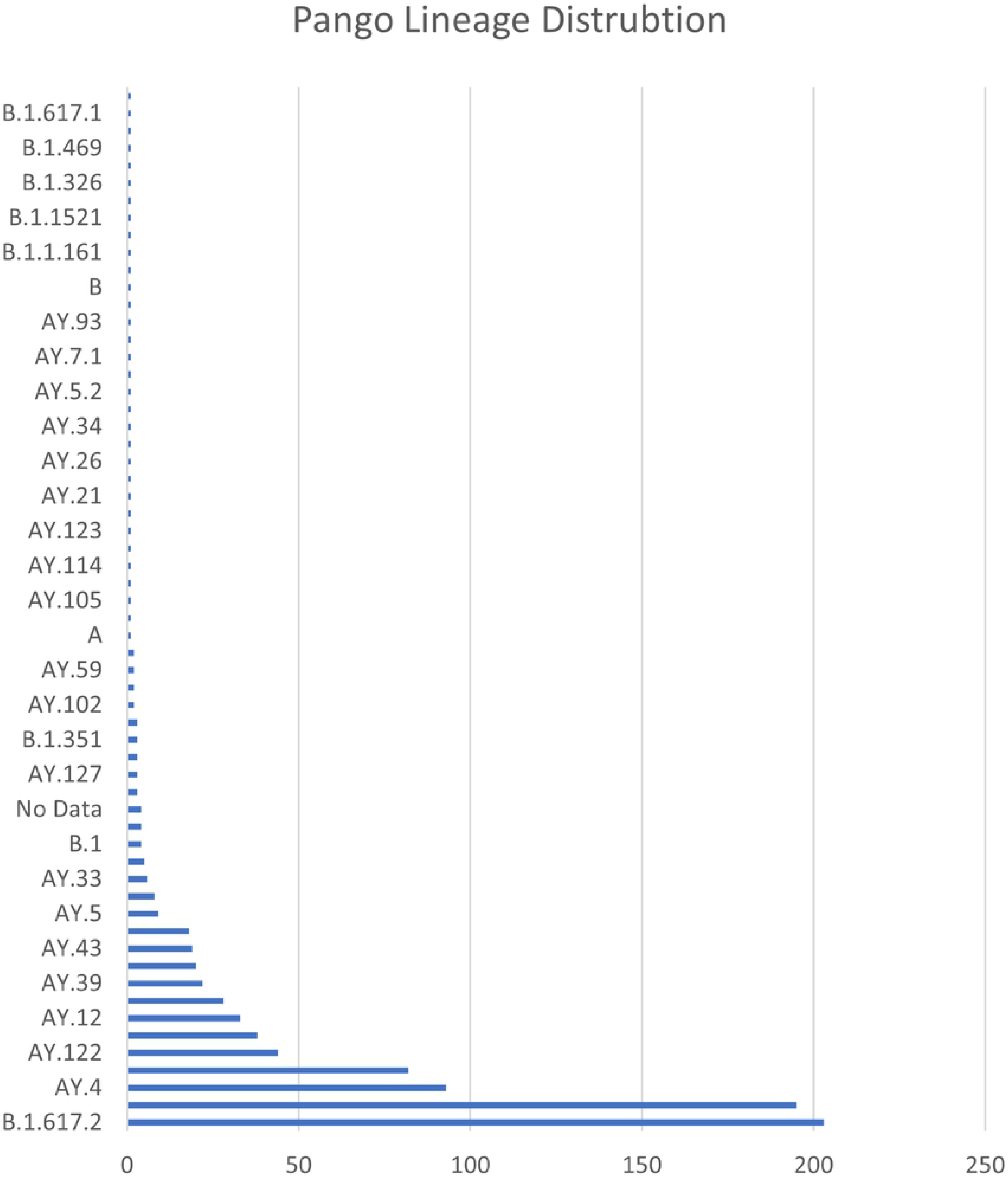
Pango Lineage Distribution of the samples in Turkey.

**Fig 3.**
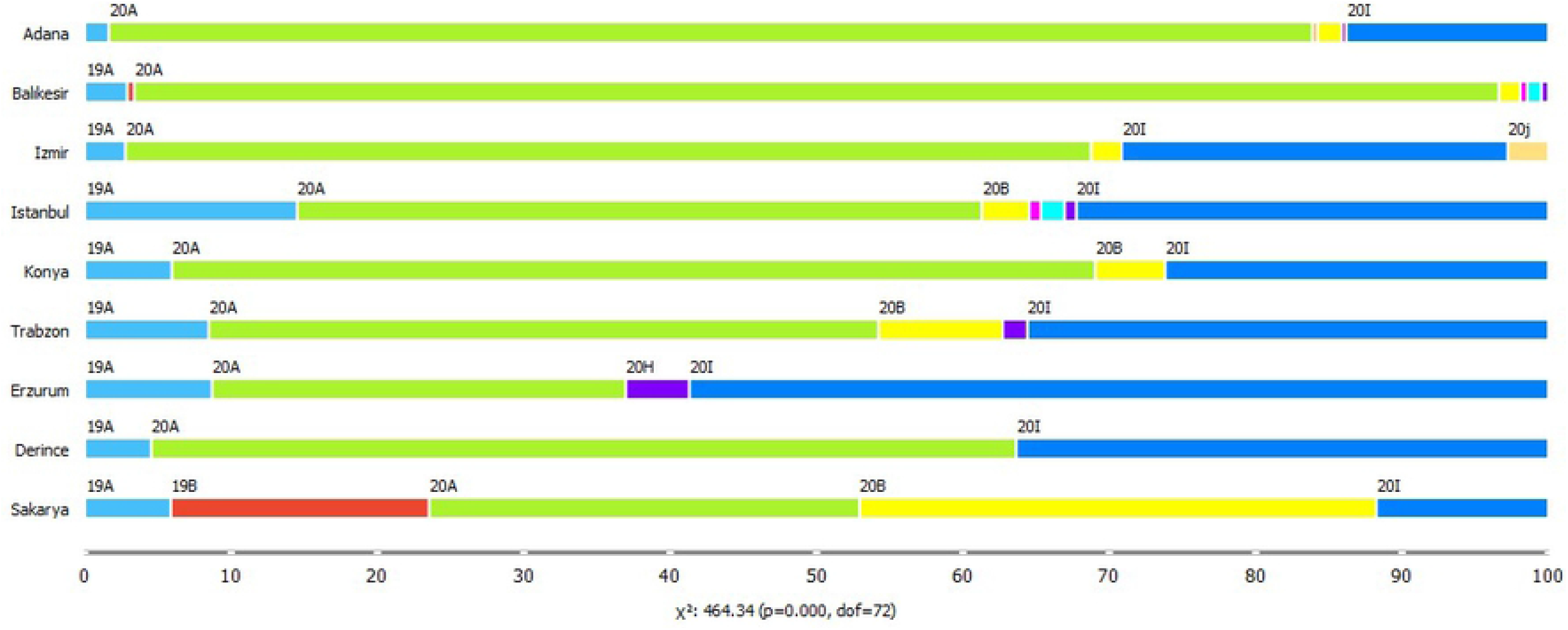
Distributions of Nextclade Clades by cities.

**Fig 4.**
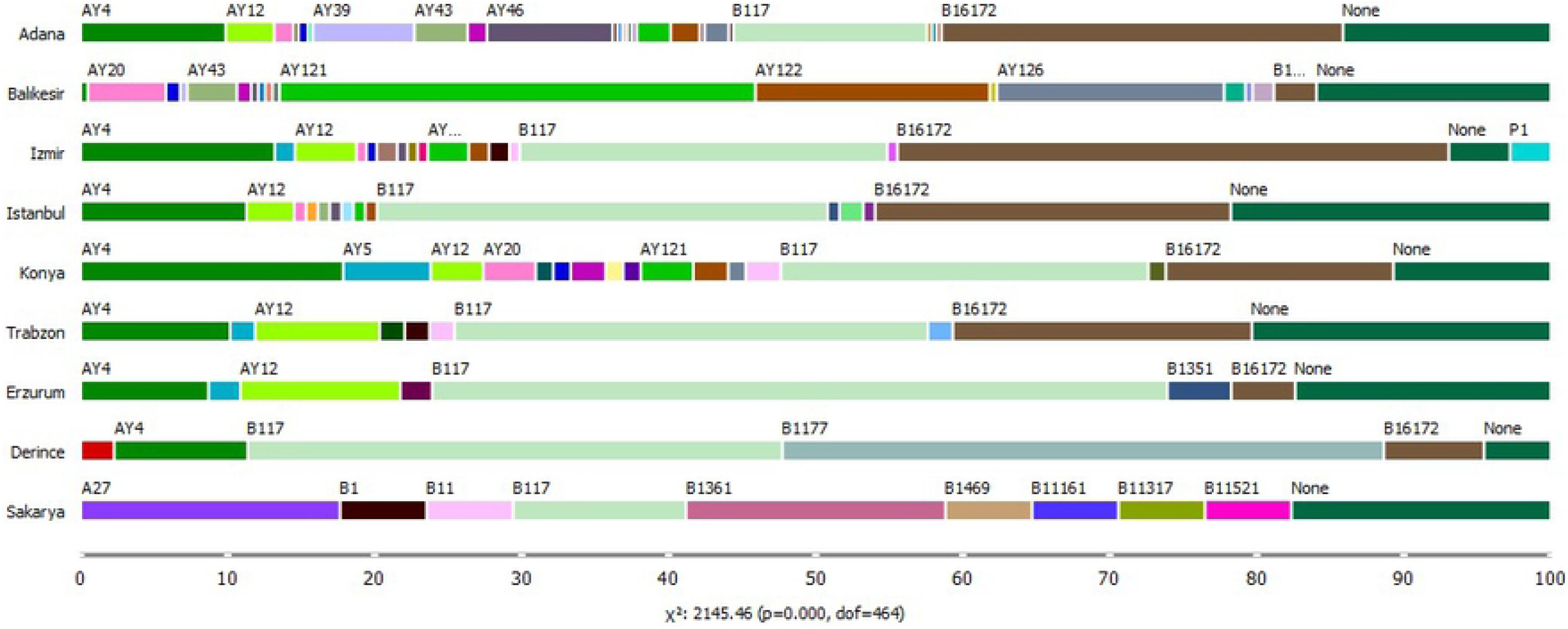
Distributions of Pango Lineages by cities.

**Fig 5.**
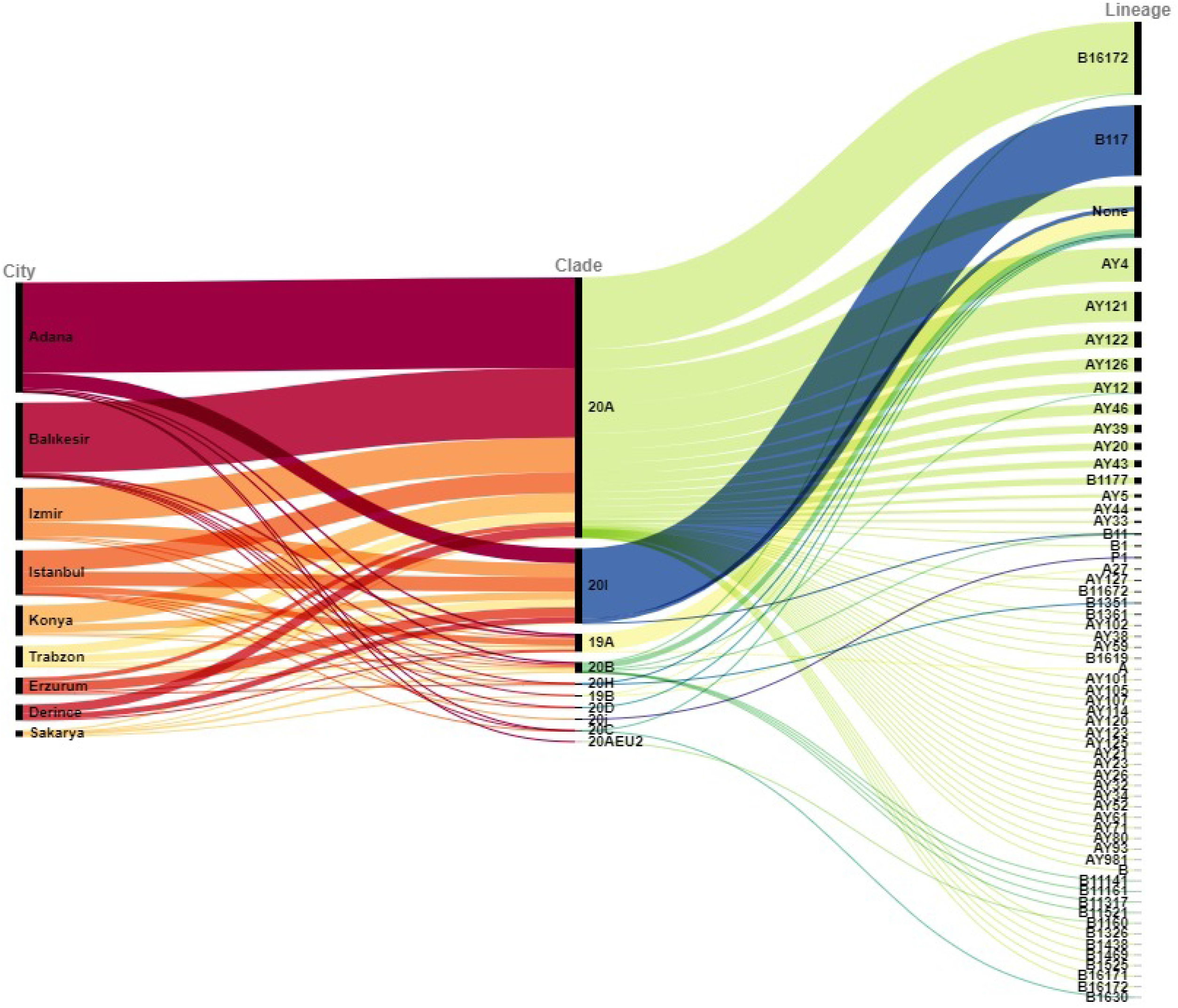
Distribution of Nexclade Clades and Pango Lineages in Turkish provinces.

**Fig 6.**
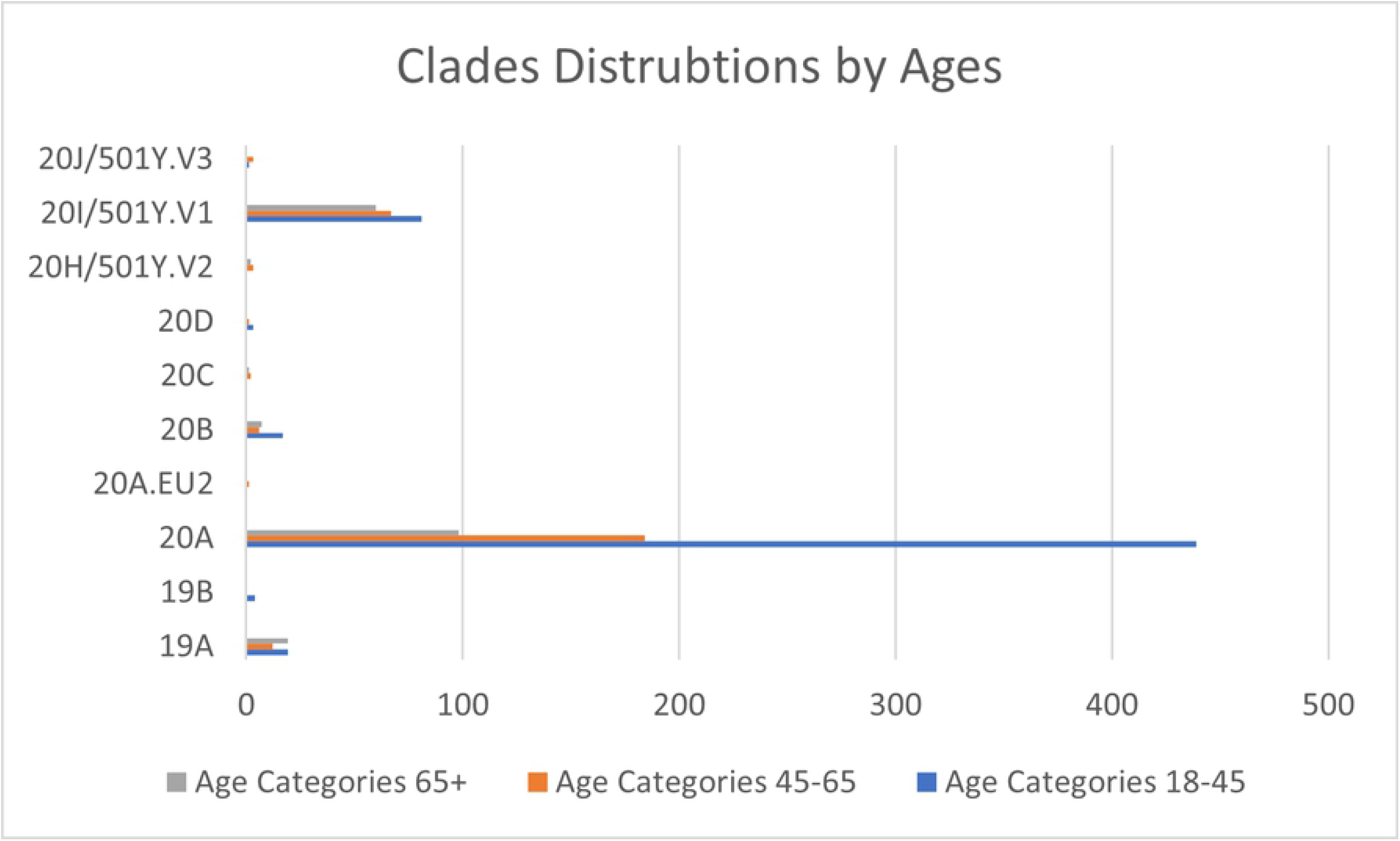
Nextclade Clade distributions by ages. The age of patients was categorized into three groups. Category 1: +65, category 2: ages between 45-65 and category 3: Ages between 18-45

**Fig 7.**
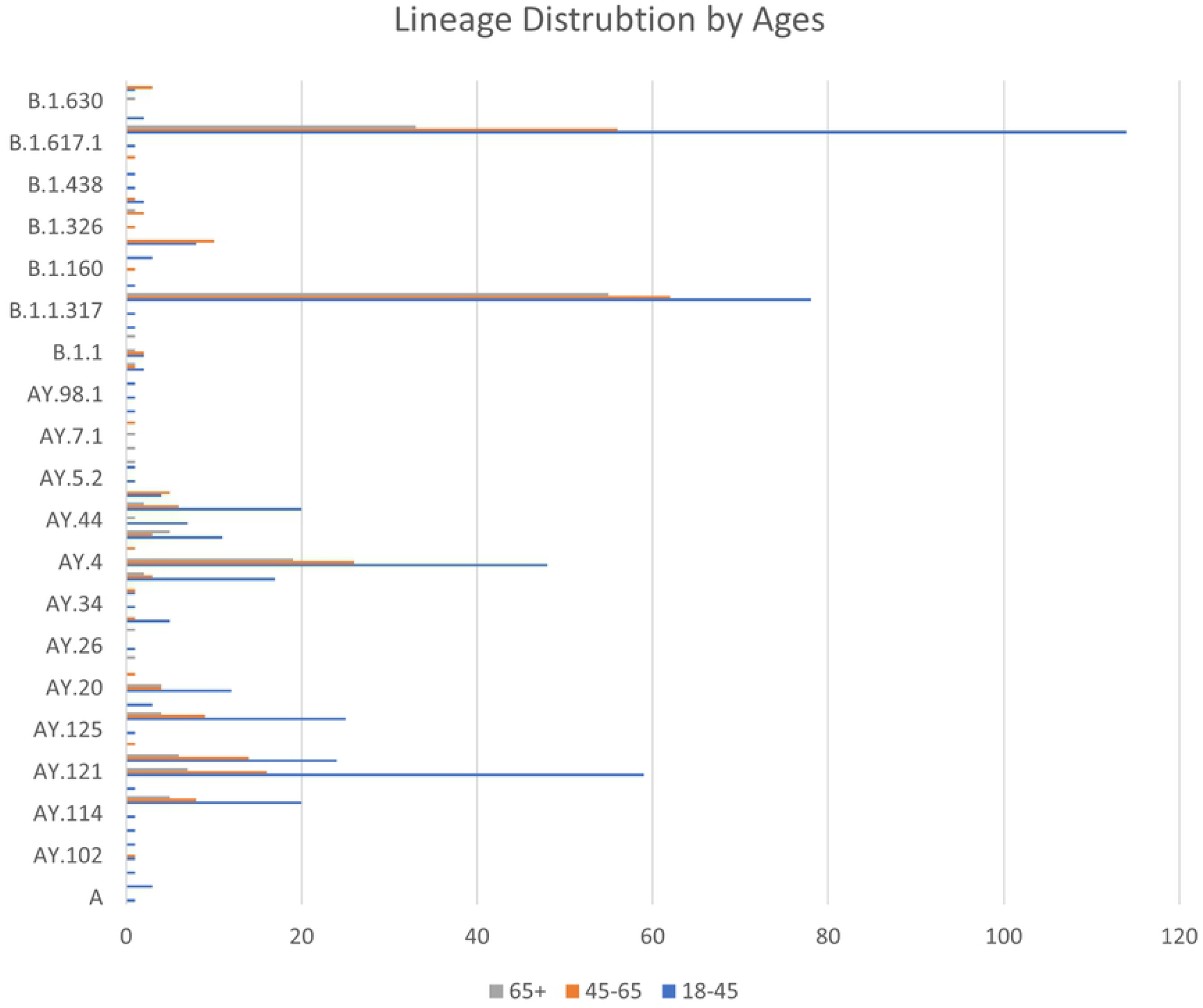
Pango Lineage distributions by ages. The age of patients was categorized into three groups. Category 1: +65, category 2: ages between 45-65 and category 3: ages between 18-45

**Fig 8.**
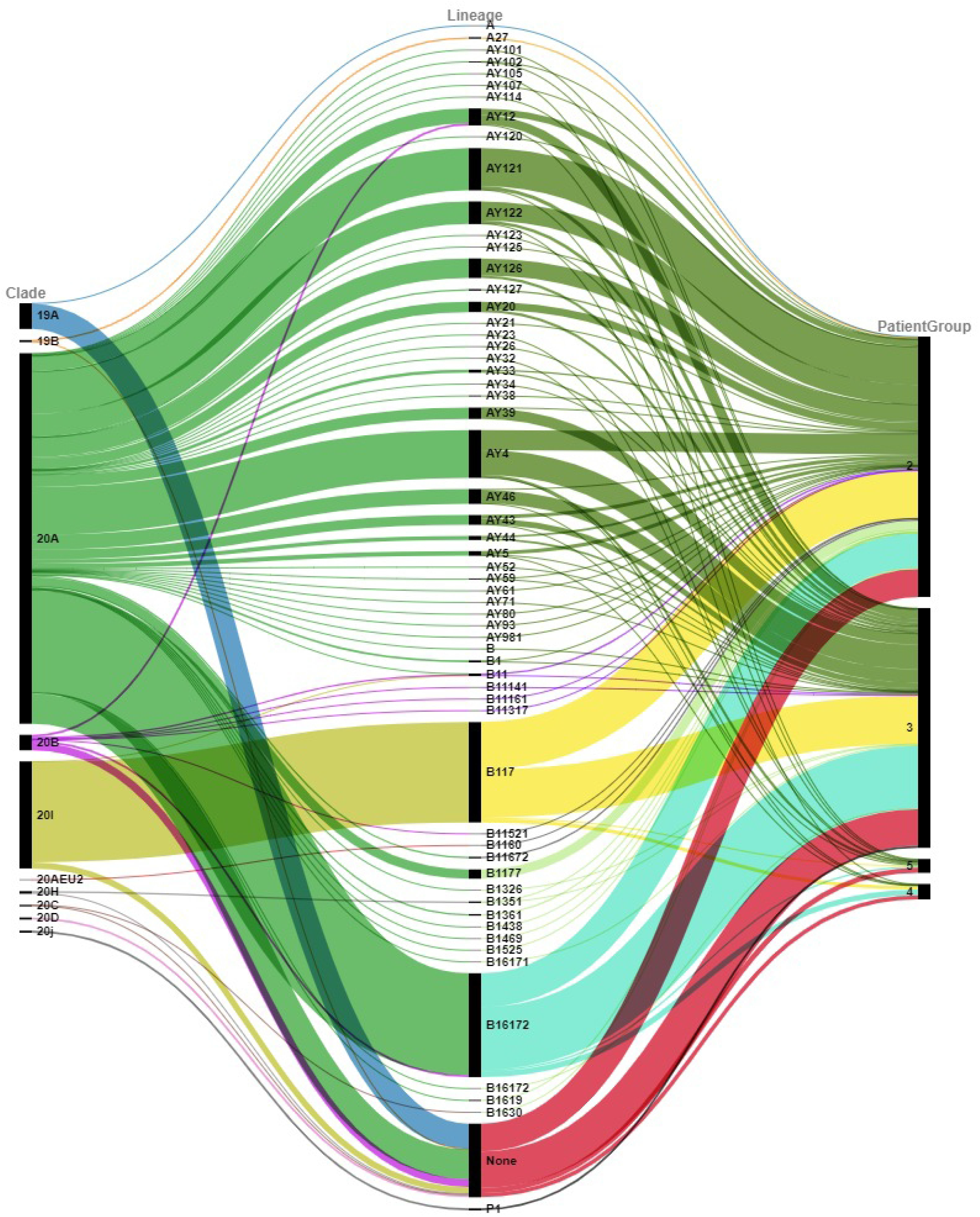
Clades and Pango Lineage distributions according to the different patient groups. Group 2: Patients with no or very mild symptoms. Group 3: patients who need hospitalization. Group 4: Patients in need of intensive care. Group 5: Patients who died in intensive care

**Fig 9.**
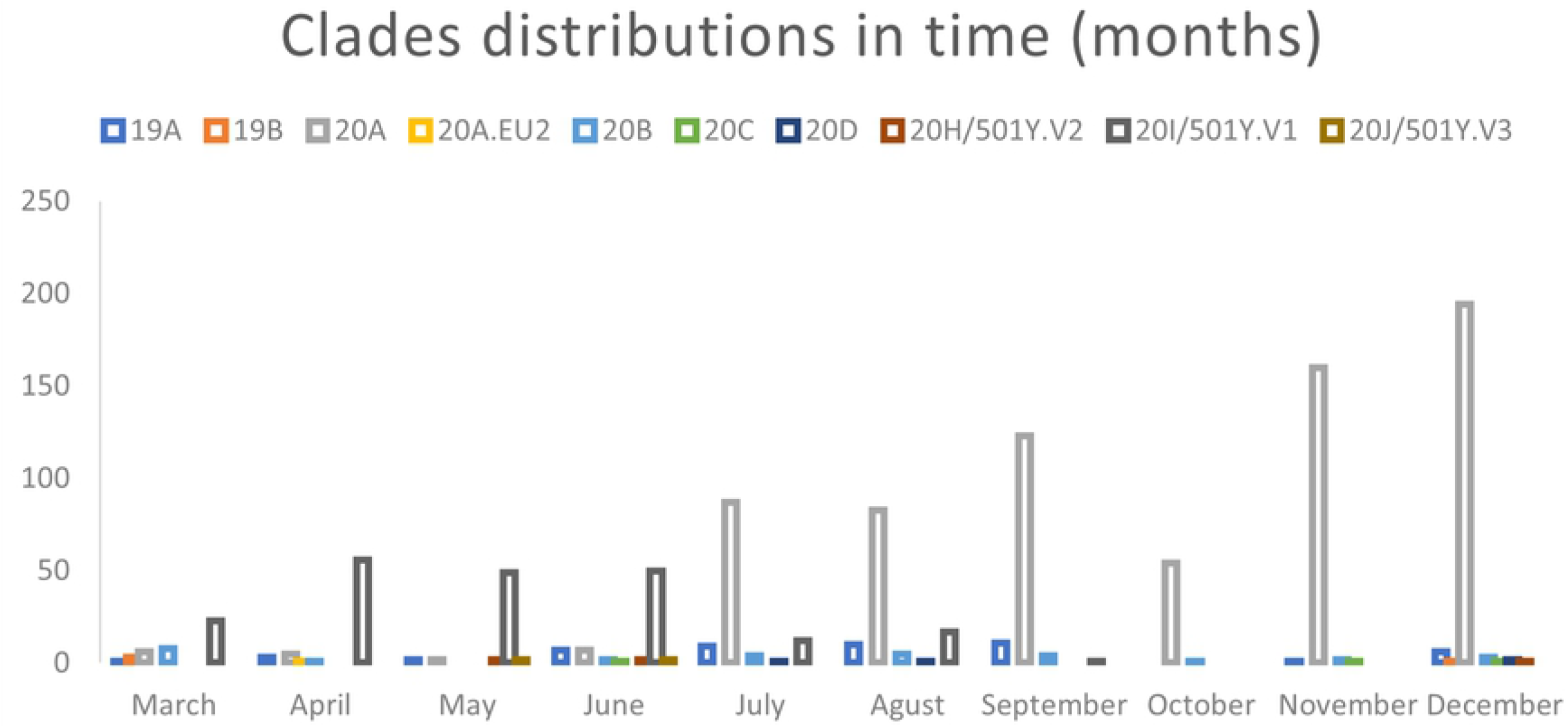
Clade distributions in each months of sample collections.

**Fig 10.**
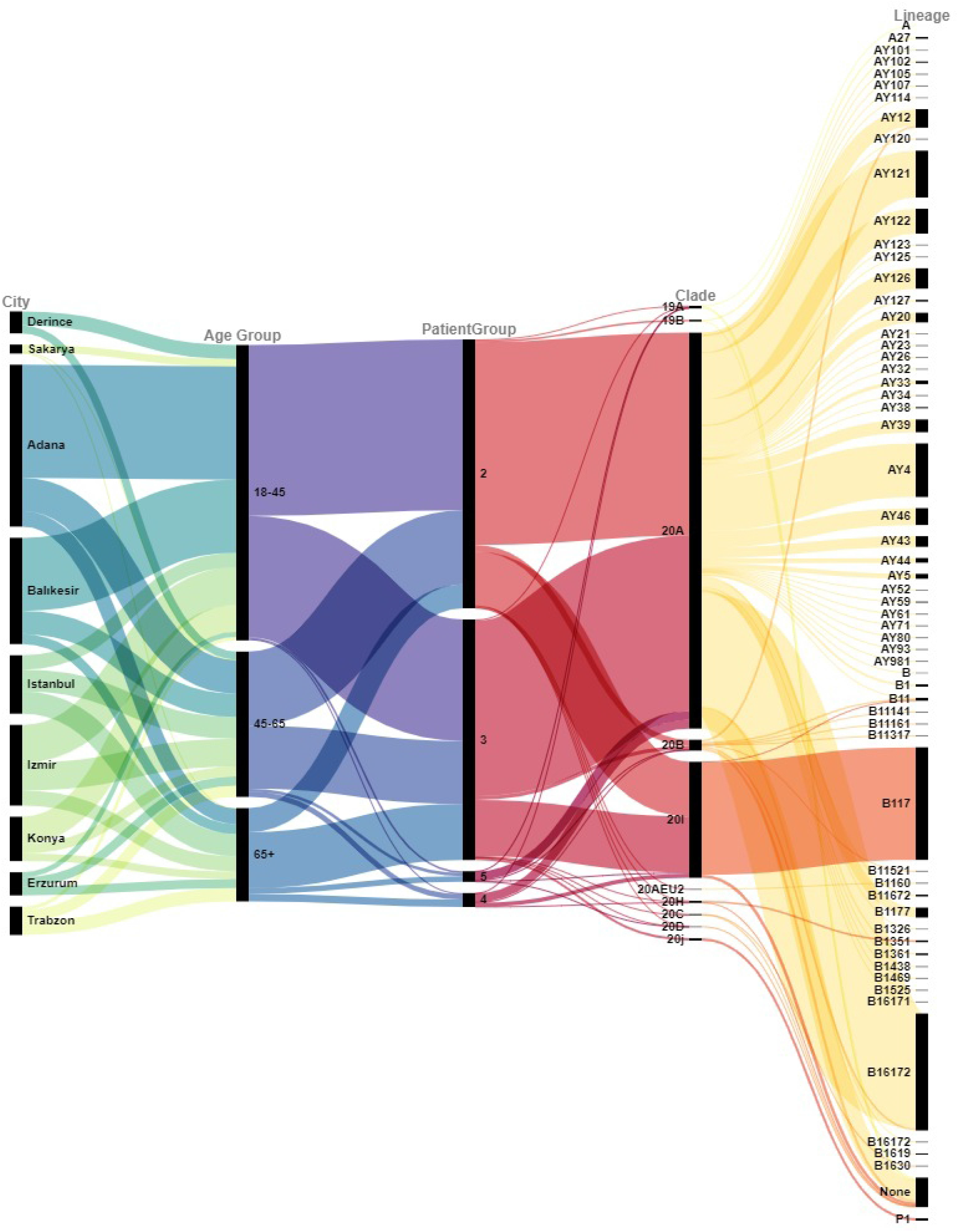
Distribution of Pango Lineages by cities, age, and patient groups.

In the following figure, we showed clades and Pango Lineage distributions among the cities together.

Since the very beginning of the pandemic, many findings have been reported that COVID-19 disease causes more damage and can be more deadly in older people (31,32). These findings have a medical explanation. Since elderly people are more likely to have chronic diseases (diabetes, cardiovascular diseases, high blood pressure, etc.), we can assume that people with these diseases are more disadvantaged when fighting the disease (33). We looked at these findings with a different approach and wanted to know if some variants of the virus are more contagious in different age groups. We grouped the ages of the patients into 3 categories: 18-45, 45-65 and over 65. Then we examined the distributions of all variants in these three age categories.

While 77% of patients between the ages of 18-45 have the 20A variant, the rate of patients over 65 years of age is slightly more than 50% in this category. The relationship between variation and age is also remarkable in the 19A clade. While 10% of patients over 65 years of age are in this clade, only 3% of patients in the 18-45 and 45-65 categories are in the 19A clade. Another interesting variation is the 20I/501Y.V1. For this clade, 35% of the patients over 65 years of age are seen in this clade, and it is 14% in the 18-45 age category and 24% in the 45-65% categories.

In this study, we also analyzed the effect of variations on disease severity. We categorized disease severity into five groups Group 1: Control group (PCR negative) Group 2: Patients with no or very mild symptoms. Group 3: patients who need hospitalization. Group 4: Patients in need of intensive care. Group 5: Patients who died in intensive care. From the moment the first variant of the virus appeared, the scientific world began to be interested in the effect of the virus on disease prognosis. Many studies on this subject have shown that various variants of the virus have various effects on the rate of transmission of the disease, the incubation period of the virus, and the severity of the disease (34–37). As seen in the table, the virus loses its lethality as the virus evolved. Patients infected with the newer variants appear to contract the disease less severely. Especially the patients in the 5th category in the 19A clade, this category consists of patients who needed intensive care and passed away, the frequency of being in 19A is much higher than the incidence of 19A in other groups. On the other hand, when 20A is examined, it is seen that the frequency of patients who do not show symptoms or who have survived the disease with very mild symptoms is much higher than the other groups.

We also tried to follow the spread of the virus variants in 10 month periods from March 2021 to December 2021.

Since the beginning of the pandemic, we have observed that the mutations seen especially in the spike protein have resulted in the emergence of different SARS-CoV-2 variants, which in turn affected the epidemiological and clinical aspects of the COVID-19 pandemic (4). Researchers have shown that variants may increase the rate of virus transmission and the risk of re-infection, and reduce the protection provided by neutralizing antibodies and vaccination (38,39). Although some countries lackof the infrastructure and qualified personnel to conduct systemic genome analysis of SARS-CoV-2, NGS tools and open access online databases of genome information make routine monitoring of mutations and variants possible (40).

Variants of SARS-CoV-2 that have spread widely and have proven to be more transmissible and have a greater impact on disease severity and immunity have been classified as Variant of Concern (VOC) by the World Health Organization (WHO). According to WHO there were five variant of concerns (VOC): Alpha B.1.1.7 (1-Dec-2020), Beta B.1.351 (18-Dec-2020), Gamma P.1 (11-Jan-2021), Delta B.1.617.2 (11-May-2021), and Omicron B.1.1.529 (26-Nov-2021) variants. Currently, only Delta and Omicron are considered as VOC (23). Although most of the variant of concern were found in our analysis, we did not detect any omicron variant in our analysis (Ministry of Health of Turkey has announced the first 6 Omicron variant on Dec 11, 2021).

Variants containing mutations similar to those found in VOCs but have spread less widely have been named as variants of interest (VOI). These have also significant effects on contagiousness, severity of disease and immunity. Some of the Previously Circulating VOIs are Lambda C.37 (14-Jun-2021), Mu B.1.621 (30-Aug-2021) and Kappa B.1.617.1 (4-Aprl-2021). Currently, there is no circulating VOI (March, 2022) (23).

According to these findings, improving genomic surveillance is critical for the understanding spread of SARS-CoV-2 in different countries. Identifying possible worldwide transmission networks quickly and consolidating response strategies is crucial to fight against to virus. In this manner, this study is important as the first extensive genomic surveillance of SARS-CoV-2 in Turkey. The real-time updating of genome sequences allows researchers to follow the virus’s most recent genetic evolution and the spread of emergent clades. The quantity of accessible sequences varies greatly between countries, and with more frequent and faster sequencing efforts across Europe and the world, we may be able to uncover new variations sooner.

In this regard, a general picture of SARS-CoV-2 variants seen in the different regions, different ages and different disease severity has summarized in the following figure.

SARS-CoV-2 continuously evolve as genetic mutations occur during replication of the genome. SARS-CoV-2 genetic lineages are routinely monitored through epidemiological investigations in some countries. Variants with specific genetic markers and variants for which there is evidence of an increase in transmissibility were tracked. Definition of variants and genomic surveillance on SARS-CoV-2 are important for better assessment of COVID-19 epidemiology. Variations and clinical patterns relations was reported in several studies (41,42). In this paper, we report the distribution of variations within 1000 SARS-CoV-2 genome samples and the expansion/progression of SARS-CoV-2 in different geographical regions of Turkey. This is the most comprehensive SARS-CoV-2 genome analysis done in Turkey during COVID-19 pandemics. The data that we have can answer the question of whether there was a unique variation in the geography that is the subject of the study, and although we saw many unique mutations in our study, these mutations need to be studied in more detail. In addition, this data will be useful in predicting new upcoming mutations in the viral genome and in determining with which variations the course of the pandemic will continue.

## Materials and Methods

### Sample Collection

Nasopharyngeal swab samples obtained from patients diagnosed with SARS-CoV-2 infection by performing a RT-PCR (Coronex, Bio-Speedy and Diagno5plex NS) in hospitals between March 2021 and December 2021 were included in the study (the approval for this research was obtained from the ethics committee of University of Health Sciences). At the beginning of the study, a website, which only could be accessed by the clinicians and researchers involved in this study, was created to ensure the coordination of the samples and the secure flow of information. 1346 nasopharyngeal swab samples were collected from nine different regions of Turkey (Fig 11). Clinicians within the researchers uploaded the information of the collected samples to the website and the recorded samples were transferred to the laboratory under suitable conditions for RNA extraction (Fig 12).

**Fig 11.**
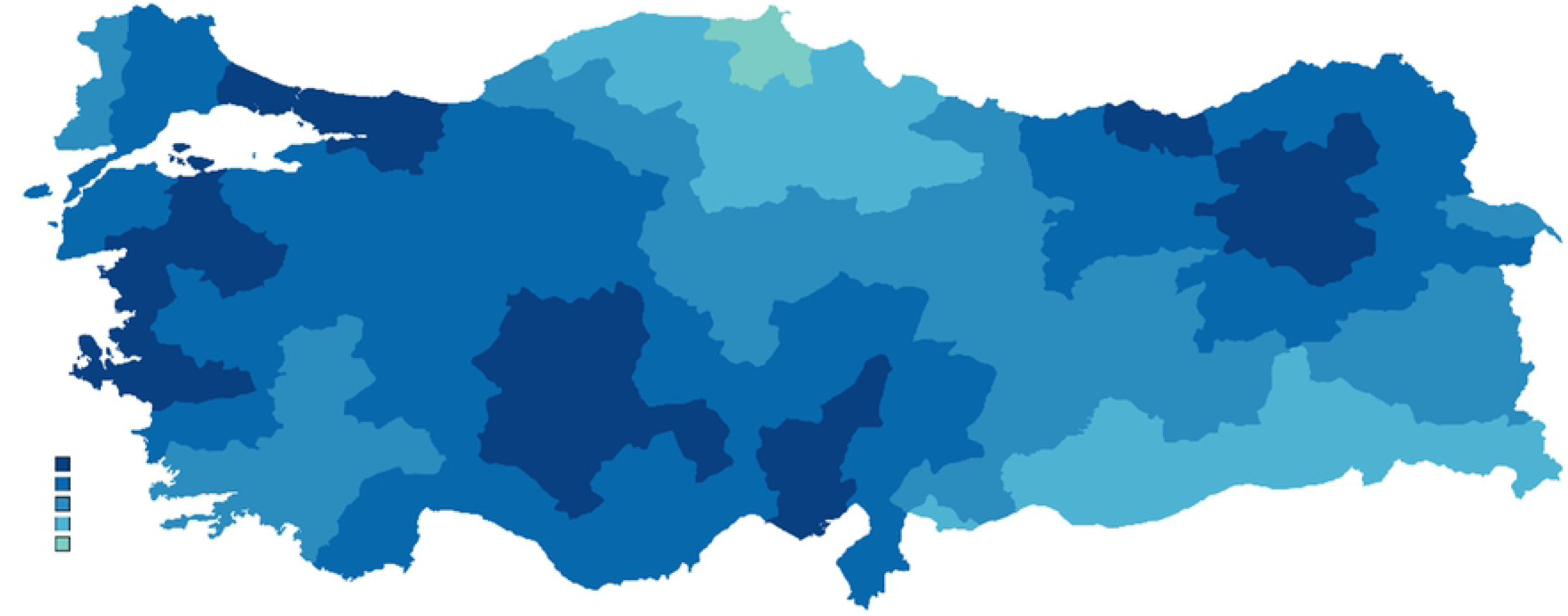
Geographic distribution of the collected SARS-CoV-2 samples. Sample collection sites are marked in dark blue. The source of sample decrease as the color becomes lighter. (Figure was drawn in https://mapchart.net/turkey.html)

**Fig 12.**
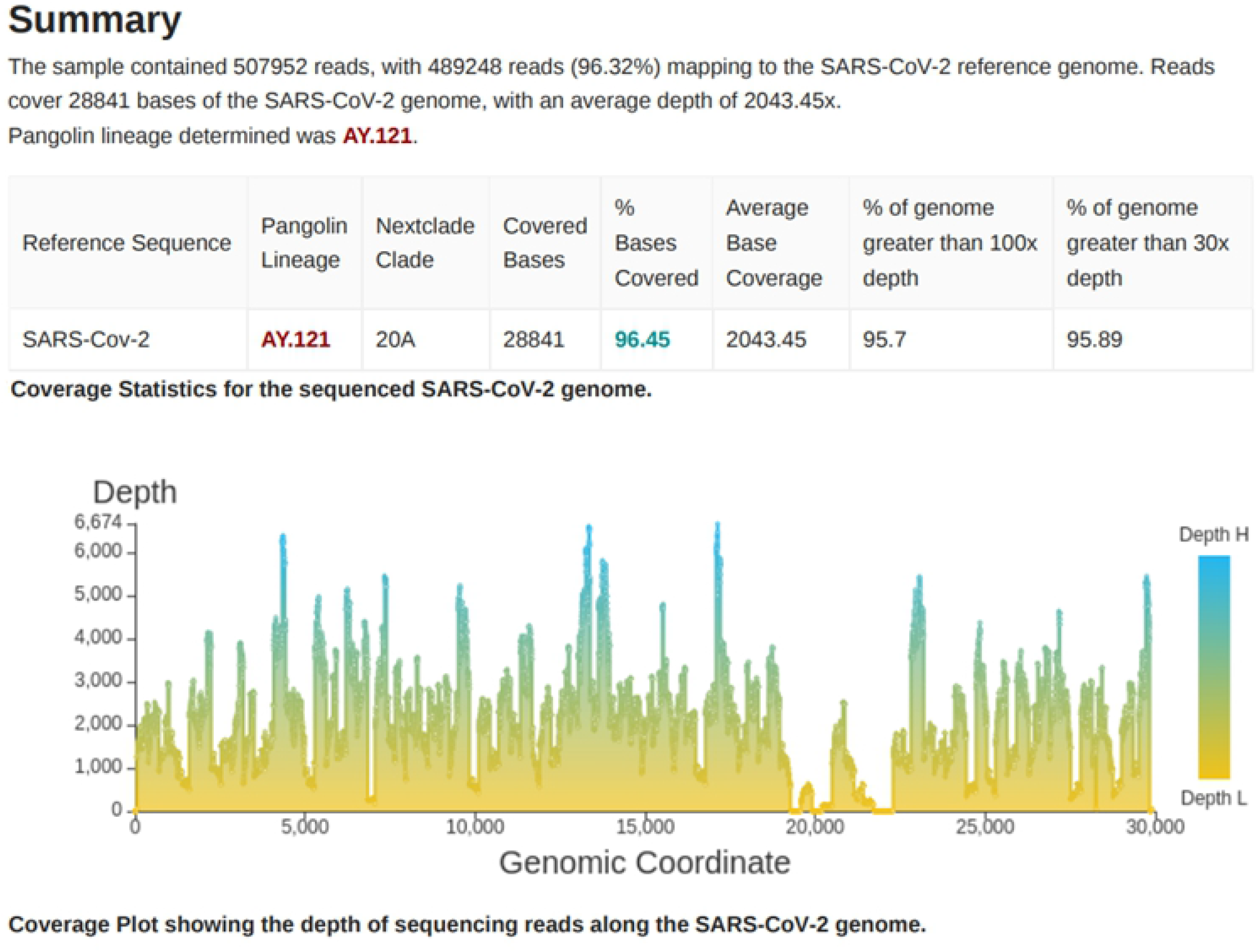
Report generated using CosmosID/the SARS-CoV-2 Strain Typing Analysis Portal includes a table indicating coverage statistics for each sequenced viral genome including information about Pango lineage and Nextclade clades and a coverage plot.

### Viral RNA Extraction

All collected samples were transferred to Genetic Engineering and Biotechnology Institute of the Scientific and Technological Research Institution of Turkey TÜBİTAK for viral RNA (vRNA). RNA isolation of 1580 samples that presented cycle threshold (CT) values between 12 and 28 were extracted using the Quick-RNA™ Viral kit (Zymo Research, USA) according to the manufacturer’s instructions in a Biosafety Level-3 laboratory. Samples with CT> 28 were not further processed.

RNA quality and quantity were evaluated with NanoDrop™ 2000 (260/280 nm and 260/230 nm ratios) and Qubit™ 3.0 Fluorometer using Qubit™ RNA HS Assay Kit (Thermo Fisher Scientific Inc., USA). The samples were stored at −80 °C until transported to the Experimental Medicine Research and Application Center of the University of Health Sciences in Istanbul for library preparation and sequencing.

### Library Preparation and SARS-CoV-2 Genome Sequencing

SARS-CoV-2 genome libraries were constructed by using the NEXTFLEX® Variant-Seq™ SARS-CoV-2 kit with the automated high-throughput liquid handler Sciclone® G3 NGS (Perkin Elmer, USA) according to the manufacturer’s instructions.

The concentration of the final libraries was measured using Qubit™ dsDNA HS Assay Kit on a Qubit™ 4.0 Fluorometer (Thermo Fisher Scientific Inc., USA) according to the manufacturer’s instructions. Quality assessment was conducted by using D1000 ScreenTape Assay and the 4150TapeStation system (Agilent Technologies, USA). Sequencing of prepared libraries was performed using the Illumina® High Output Kit and MiniSeq™ platform according to manufacturer’s protocol with paired-ends reads (2×149 bp). 1.4 pM of the pooled library with 1% phiX was loaded.

### *In silico* Characterization of SARS-CoV-2 Genome

The raw data was obtained in BCL format and converted to the FASTQ format using Bcltofastq X Terminal. Then, those FASTQ files were uploaded to the SARS-CoV-2 Strain Typing Analysis Portal (CosmosID®). By using this platform reads were mapped to the SARS-CoV-2 reference genome (NC_045512). Accurate information about SARS-CoV-2 genome coverage was obtained for each of the samples, together with Pangolin lineages and Nextclade clades. Correspondence of a detected lineage to a variant of concern was also indicated. An example of the analysis report of this pipeline is shown in Figure 12.

The SARS-CoV-2 Strain Typing Analysis Portal detects, first, SARS-CoV-2 at the species level. Then, the platform compares Next Generation Sequencing (NGS) reads to sequence signatures (k-mers) in a database. This database is arranged as a phylogenetic tree and contains unique and shared k-mers, in this way platform is able to map each level in the tree. This unique phylogenetic structure of the database allowed us to perform a unique classification of pathogens and even detect mutations that are still unknown to the world.

